# Dysregulation of Nitrosylation Dynamics Promotes Nitrosative Stress and Contributes to Cardiometabolic Heart Failure with Preserved Ejection Fraction

**DOI:** 10.1101/2024.12.20.629549

**Authors:** Zhen Li, Kyle B. LaPenna, Natalie D. Gehred, Xiaoman Yu, W.H. Wilson Tang, Jake E. Doiron, Huijing Xia, Jingshu Chen, Ian H. Driver, Frank B. Sachse, Naoto Muraoka, Antonia Katsouda, Paraskevas Zampas, Amelia G. Haydel, Heather Quiriarte, Alexia Zagouras, Jennifer Wilcox, Tatiana Gromova, Sanjiv J. Shah, Traci T. Goodchild, Timothy D. Allerton, Martin B. Jensen, Andreas Papapetropoulos, Thomas E. Sharp, Thomas M. Vondriska, David J. Lefer

## Abstract

**Background:** Recent reports suggest increased myocardial iNOS expression leads to excessive protein *s*-nitrosylation, contributing to the pathophysiology of HFpEF. However, the relationship between NO bioavailability, dynamic regulation of protein *s*-nitrosylation by trans- and de-nitrosylases, and HFpEF pathophysiology has not been elucidated. Here, we provide novel insights into the delicate interplay between NO bioavailability and protein *s*-nitrosylation in HFpEF.

**Methods:** Plasma nitrite, nitrosothiols (RsNO), and 3-nitrotyrosine (3-NT) were measured in HFpEF patients and in controls. Studies in WKY or ZSF1 obese rats were performed to evaluate HFpEF severity, NO signaling, and total nitroso-species (Rx(s)NO) levels. snRNA sequencing was performed to identify key genes involved in NO signaling and *s*-nitrosylation regulation.

**Results:** In HFpEF patients, circulating RsNO and 3-NT were significantly elevated while nitrite, a biomarker for NO bioavailability, remained unchanged. In ZSF1 obese rats, NO bioavailability was significantly reduced while Rx(s)NO levels exhibited an age-dependent increase as HFpEF progressed. snRNA seq highlighted significant upregulation of a trans-nitrosylase, hemoglobin-beta subunit (HBb), which was corroborated in human HFpEF hearts^1^. Subsequent experiments confirmed HBb upregulation and revealed significant reductions in enzyme activity of two major de-nitrosylases, Trx2 and GSNOR in ZSF1 obese hearts. Further, elevated RxNO levels, increased HBb expression, and reduced activity of Trx2 and GSNOR were identified in the kidney and liver of the ZSF1 obese rats.

**Conclusions:** Our data reveal circulating markers of nitrosative stress (RsNO and 3-NT) are significantly elevated in HFpEF patients. Data from the ZSF1 obese rat model mirror the results from HFpEF patients and reveal that pathological accumulation of RxNO/nitrosative stress in HFpEF may be in part, due to the upregulation of the trans-nitrosylase, HBb, and impaired activity of the de-nitrosylases, Trx2 and GSNOR. Our data suggest that dysregulated protein nitrosylation dynamics in the heart, liver, and kidney contribute to the pathogenesis of cardiometabolic HFpEF.

**Translational Perspective:** Our findings describe for the first time that circulating RsNO and 3-NT are significantly upregulated in HFpEF patients suggesting systemic nitrosative stress in HFpEF, and demonstrate a profound disconnect between insufficient physiological NO signaling and pathological nitrosative stress in HFpEF, which is in stark contrast to HFrEF in which both NO bioavailability and protein *s*-nitrosylation are attenuated. Further, this study provides novel mechanistic insights into a critical molecular feature of HFpEF in humans and animal models: nitrosative stress arises predominantly from imbalance of trans-nitrosylases and de-nitrosylases, thereby leading to impaired NO bioavailability concomitant with increased protein *s*-nitrosylation. Importantly, these perturbations extend beyond the heart to the kidney and liver, suggesting HFpEF is characterized by a systemic derangement in trans- and de-nitrosylase activity and providing a unifying molecular lesion for the systemic presentation of HFpEF pathophysiology. These findings have direct clinical implications for the modulation of NO levels in the HFpEF patient, and indicate that restoring the balance between trans- and denitrosylases may be novel therapeutic targets to ameliorate disease symptoms in HFpEF patients.

## INTRODUCTION

Heart failure with preserved ejection fraction (HFpEF) is among the most complex cardiovascular diseases. Over the last decade, the prevalence of HFpEF has surpassed heart failure with reduced ejection fraction (HFrEF) and has become the most common form of heart failure in the United States^2–5^. HFpEF patients exhibit a combination of comorbidities including obesity, diabetes, liver and renal dysfunction, aging, and hypertension. Due to its heterogeneous nature and lack of effective therapies, clinical management of HFpEF is exceedingly challenging. Additionally, despite intense efforts in recent years, the pathophysiological, cellular, and molecular factors that contribute to HFpEF have yet to be precisely identified^2^. As a result, very few therapies have achieved clinical efficacy, with SGLT2 inhibitors, angiotensin receptor neprilysin inhibitor (i.e, LCZ696), and mineralocorticoid receptor antagonist (i.e, spironolactone) as the only pharmacological agents approved by the FDA for the use in HFpEF^6–11^ while glucagon like peptide 1 (GLP-1) - based therapies have demonstrated benefits in clinical trials involving HFpEF patients with diabetes and obesity. Yet, no treatment regimen (medication or device) has been shown to reduce all-cause mortality and cardiovascular mortality in HFpEF. Discovery of novel drug targets and development of new therapeutics for the treatment of HFpEF are urgently needed.

Nitric oxide (NO) is an endogenously produced gaseous signaling molecule crucial to cardiovascular homeostasis. In heart failure with preserved ejection fraction (HFpEF), as with many other cardiovascular diseases, NO availability and signaling are impaired^12–16^; however, therapies aimed either to replenish NO bioavailability (i.e, inhaled nitrite and isosorbide mononitrate) or to facilitate NO downstream signaling (i.e, sGC stimulators and activators, PDE inhibitors) have yielded disappointing results^17–21^. Preclinical studies in rodent models of HFpEF reported that myocardial nitrosative stress, characterized by elevated total nitroso - species (RxNO), which result from overproduction of NO by inducible NO synthase (iNOS) or neuronal NO synthase (nNOS) plays a causative role in the pathogenesis and progression of HFpEF^22, 23^.

*S*-nitrosylation is a protein post-translational modification (PTM) characterized by the reversible, covalent addition of a NO moiety to the thiol group of the cysteine residues of proteins. Such PTM is considered a vital process for NO to elicit its bioactivity. While NO generated via NOS activity within cells contributes to protein *s*-nitrosylation via reaction of cysteine thiols with NO-derived metabolites, such as nitrite and N_2_O_3_, this PTM is delicately regulated by the dynamic addition and removal of nitroso-groups from cystine residues through enzymatic reactions^24, 25^. More specifically, trans-nitrosylases mediate the addition of NO moiety onto the cysteine residue while de-nitrosylases remove the NO moiety to effectively regulate the extent of protein *s*-nitrosylation and maintain physiological processes^24, 25^. Common trans-nitrosylases include hemoglobin, glyceraldehyde 3-phosphate dehydrogenase (GAPDH), and caspase-3 while common de-nitrosylases consist of S-nitrosoglutathione reductase (GSNOR), thioredoxin, and alpha-keto reductase family 1 - member 1 (AKR1a1) ^24, 26–28^. Impaired function of *s*-nitrosylation by the trans- and de-nitrosylase enzymes contributes to a broad spectrum of human diseases including but not limited to neurodegenerative disease, Duchenne muscular dystrophy, type-2 diabetes, myocardial infarction, pulmonary arterial hypertension, and pre-eclampsia^27–37^. However, the roles of trans- and de-nitrosylases have not been elucidated in the setting of HFpEF, despite their potential involvement in excessive protein *s*-nitrosylation that has previously been reported^22, 23^.

In the present study, we sought to verify the previous findings of excessive cardiac nitrosative stress and accumulated protein *s*-nitrosylation in clinical HFpEF. Moreover, we studied the regulation of trans- and de-nitrosylases to improve our understanding of the regulation of *s*-nitrosylation in HFpEF using two well-established rodent models of HFpEF, the ZSF1 obese rat model and the “two-Hit” mouse model^38–40^.

## MATERIALS AND METHODS

### Human Plasma Samples

Plasma samples were collected from either ambulatory patients with HFpEF prospectively enrolled in the outpatient clinic (with symptoms of heart failure and echocardiogram showing left ventricular ejection fraction ≥50%) or age- and sex-matched control subjects recruited from the community without existing cardiovascular diseases (confirmed by echocardiography, pulmonary function testing, and cardiac biomarkers) under protocols approved by the Institutional Review Board (IRB# 06-805 and #10-727, respectively) at the Cleveland Clinic. All participants were provided with written informed consent. Baseline characteristics of all participants are shown in ***Table 1***.

**Table 1.**
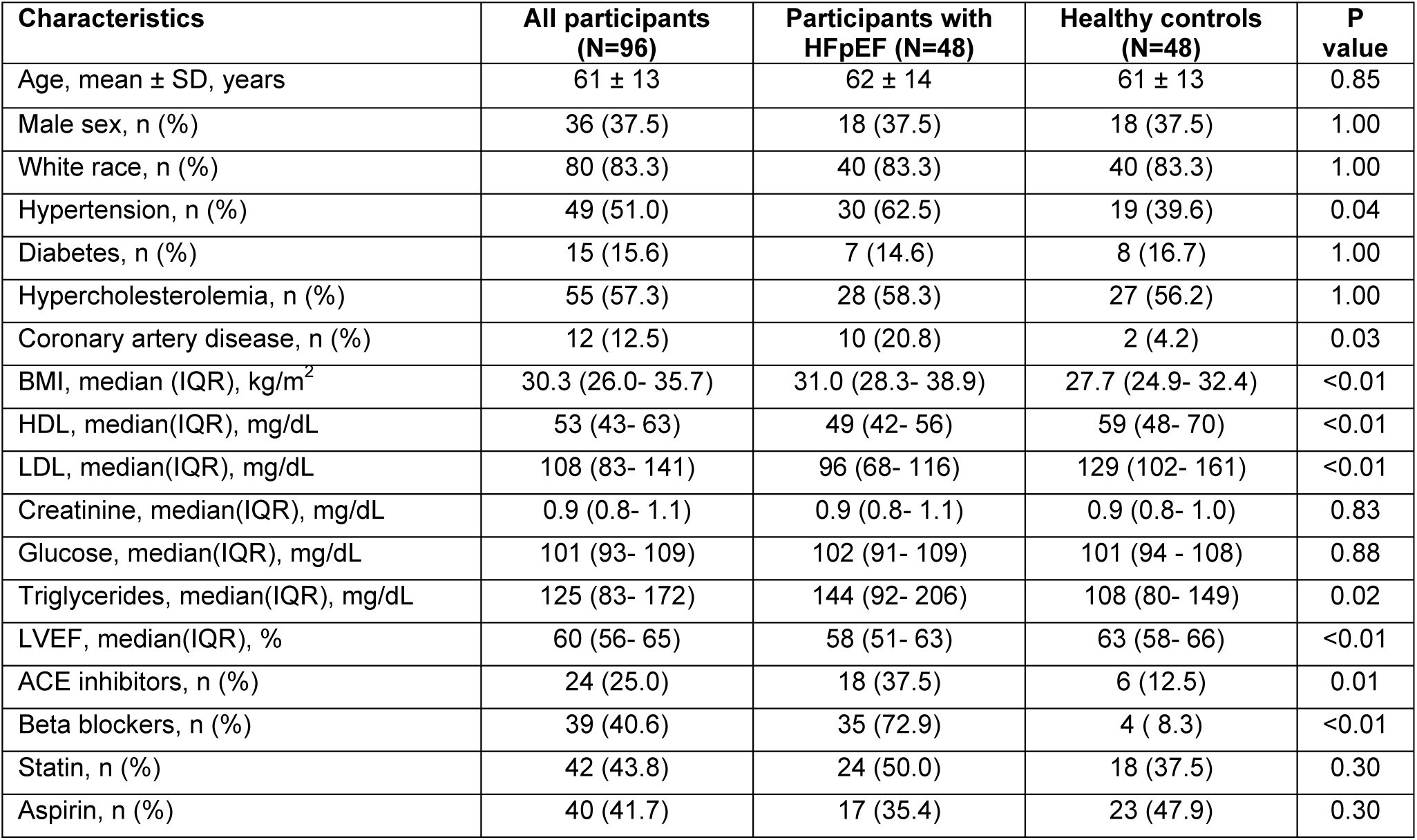
Baseline Characteristics of participants with HFpEF vs. healthy controls. The Wilcoxon–rank sum test or Welch two sample t-test for continuous variables and the χ2 test or Fisher’s exact test for categorical variables were used to determine significant difference between groups.

| Characteristics | All participants<br>(N=96) | Participants with<br>HFpEF (N=48) | Healthy controls<br>(N=48) | P<br>value |
| --- | --- | --- | --- | --- |
| Age, mean ± SD, years | 61 ± 13 | 62 ± 14 | 61 ± 13 | 0.85 |
| Male sex, n (%) | 36 (37.5) | 18 (37.5) | 18 (37.5) | 1.00 |
| White race, n (%) | 80 (83.3) | 40 (83.3) | 40 (83.3) | 1.00 |
| Hypertension, n (%) | 49 (51.0) | 30 (62.5) | 19 (39.6) | 0.04 |
| Diabetes, n (%) | 15 (15.6) | 7 (14.6) | 8 (16.7) | 1.00 |
| Hypercholesterolemia, n (%) | 55 (57.3) | 28 (58.3) | 27 (56.2) | 1.00 |
| Coronary artery disease, n (%) | 12 (12.5) | 10 (20.8) | 2 (4.2) | 0.03 |
| BMI, median (IQR), kg/m<sup>2</sup> | 30.3 (26.0- 35.7) | 31.0 (28.3- 38.9) | 27.7 (24.9- 32.4) | <0.01 |
| HDL, median(IQR), mg/dL | 53 (43- 63) | 49 (42- 56) | 59 (48- 70) | <0.01 |
| LDL, median(IQR), mg/dL | 108 (83- 141) | 96 (68- 116) | 129 (102- 161) | <0.01 |
| Creatinine, median(IQR), mg/dL | 0.9 (0.8- 1.1) | 0.9 (0.8- 1.1) | 0.9 (0.8- 1.0) | 0.83 |
| Glucose, median(IQR), mg/dL | 101 (93- 109) | 102 (91- 109) | 101 (94 - 108) | 0.88 |
| Triglycerides, median(IQR), mg/dL | 125 (83- 172) | 144 (92- 206) | 108 (80- 149) | 0.02 |
| LVEF, median(IQR), % | 60 (56- 65) | 58 (51- 63) | 63 (58- 66) | <0.01 |
| ACE inhibitors, n (%) | 24 (25.0) | 18 (37.5) | 6 (12.5) | 0.01 |
| Beta blockers, n (%) | 39 (40.6) | 35 (72.9) | 4 ( 8.3) | <0.01 |
| Statin, n (%) | 42 (43.8) | 24 (50.0) | 18 (37.5) | 0.30 |
| Aspirin, n (%) | 40 (41.7) | 17 (35.4) | 23 (47.9) | 0.30 |

### Experimental Animal Models of HFpEF

#### ZSF1 Obese Rat Model of Cardiometabolic HFpEF

Both male Wistar Kyoto rats and ZSF1 obese (Ob) rats (Charles River Laboratories, Wilmington, MA) were used in the present study. ZSF1 Ob rats spontaneously developed severe cardiometabolic HFpEF as previously charaterized^38, 39, 41–46^. Both the ZSF1 obese rat and WKY were investigated at two separate timepoints: 14 and 26 weeks of age. Group sizes of n = 6 to 8 were determined using a power and sample analysis with the significance level at 5% and power at 80%. Rats were housed at AALAC accredited animal facility of Cedars-Sinai Medical Center or LSUHSC in a temperature controlled and 12-hour light/dark cycle.

#### “Two-hit” Mouse Model of HFpEF

Starting at the age of 9 weeks, male C57BL/6N (Charles River Laboratories, Wilmington, MA) were treated with either L-N^G^-nitro arginine methyl ester (L-NAME) in drinking water (0.5 gram/L) and high fat diet to induce HFpEF or normal drinking water and normal diet for 10 weeks^22, 47^. Group sizes of n = 9-11 were determined using a power and sample analysis with the significance level at 5% and power at 80%. Mice were housed at AALAC accredited animal facility of Cedars-Sinai Medical Center or LSUHSC in a temperature controlled and 12-hour light/dark cycle.

Only male rats and mice were utilized in the current manuscript as female HFpEF patients are more likely to display lower body mass index as compared to their male counterpart^48^, which differs from the cardiometabolic phenotypes of the ZSF1 Ob rat and “two-Hit” mouse models of HFpEF. Additionally, previous studies have highlighted the cardiovascular protective effects of estrogen in female sex^49, 50^, thereby studies using ovariectomized or postmenopausal aging female rodents are warranted in the future to fully investigate the dynamic controls of protein *s*-nitrosylation/nitrosative stress in HFpEF for both sexes.

All studies were Cedars-Sinai Medical Center or LSUHSC IACUC (Institutional Animal Care and Use Committee) approved and received care according to AALAC guidelines.

### Nitrite Measurements

Both plasma and tissue homogenate nitrite quantification were performed using a high-performance liquid chromatography system (ENO30 Analyzer, Amuza Inc., San Diego, USA) specifically designed for nitrite determination as described previously^22, 51–53^. Briefly, biological fluid samples such as plasma or tissue homogenates were subjected to methanol precipitation to remove lipids and proteins. Then the supernatants were analyzed for nitrite content. Varying concentrations of pure sodium nitrite solutions between 0.039 to 5 µM were utilized to generate the standard curve for the quantification of the nitrite concentration in the samples.

### Nitroso - Species Measurements

The levels of total nitroso - species (RsNO in the plasma and RxNO in the tissue homogenates) were quantified using an ion chromatography device coupled with gas-phase chemiluminescence (nCLD88, Echo Physics, Duernten, Switzerland) as previously described^15^. Briefly, biological fluid samples such as plasma or tissue homogenates were incubated with 2.0% mercuric chloride followed by acidified sulfanilamide, then the reaction mixture were subjected to a reductive denitrosation reaction by iodine - iodide. The denitrosation reaction detaches the nitroso - group from the nitroso - species, which then is released as gas-phase NO with a one-to-one ratio. Finally, the amount of gas-phase NO was analyzed through the nCLD88 system.

### Exercise Capacity Testing

Treadmill exercise capacity of ZSF1 obese or WKY rats as well as control or HFpEF mice were measured using an IITC Life Science 800 Series rodent specific treadmill (IITC Life Science, Woodland Hills, CA). Rats first were allowed to acclimate to the treadmill environment for a period of 5 minutes. Following acclimation, rats were carried through a warm-up phase of initially 6 meters per minute with gradual increase to 12 meters per minute during a 4-minute ramp up time for a total of 5 minutes. The animals were then running at a rate of 12 meters per minute with no ramp incline until a state of exhaustion. Both duration and calculated distance were plotted along with work (kg * meter).

### Systemic and Left Ventricular Hemodynamic Measurements

At each timepoint, animals were anesthetized using isoflurane at 3-4% for induction and 2 to 2.5% for maintenance during the procedure. The right common carotid was canulated with a high-fidelity pressure catheter (Transonic, NY, US) which measured the systemic pressures at systole and diastole. The pressure catheter was then introduced into the left ventricle of the heart. Left ventricular end diastolic pressures and ventricular relaxation time constant was measured as previously described^54, 55^.

### Enzyme-linked Immunosorbent Assays (3-NT and cGMP)

The levels of 3-Nitrotyrosine were measured with commercially available ELISA assay kits in plasma samples from control or HFpEF patients (#STA-305, Cell Biolabs, San Diego, CA), and from control or HFpEF rodent models (#NBP2-66363, Novus Biologicals, Toronto, Canada). Circulating cGMP levels in plasma samples from control or ZSF1 Ob rats were measured with commercially available ELISA kit (#581021, Cayman Chemical, Ann Arbor, MI).

### Single Nuclei RNA Sequencing

Left ventricle of the heart tissue were collected from WKY and ZSF1 Obese Rats at 14 or 26 weeks of age and immediately snap-frozen in liquid nitrogen. Heart nuclei were isolated using a lysis buffer consisting of 0.25M sucrose, 10mM Tris-Hcl pH 7.5, 25mM KCl, 5mM MgCl2, 45uM Actinomycin D, supplemented with 1X protease inhibitor (G6521, Promega), 0.4U/uL RNasin Ribonuclease Inhibitor (N2515, Promega), 0.2U/ul SuperaseIn (AM2694, ThermoFisher). Briefly, heart tissue samples were minced into smaller pieces with scissors in a 1ml lysis buffer. The minced tissue was then homogenized in a Dounce homogenizer on ice with 10 strokes of pestle A, followed by 10 strokes of pestle B. The homogenates were then filtered through a 40 μm cell strainer and centrifuged at 400 x g for 5 min at 4°C. The nuclei pellet was resuspended in 2% BSA in PBS supplemented with Protector RNase inhibitor (03335402001, Sigma-Aldrich, Burlington MA) at 0.2 U/ul. Heart nuclei were stained with Sytox red (S34859, Thermo Fisher, Waltham, MA) and sorted and purified through fluorescence-activated cell sorting (FACS). 8,000-10,000 nuclei from each rat heart were processed using a 10X Genomics microfluidics chip to generate barcoded Gel Bead-In Emulsions according to manufacturers’ protocols. Indexed single-cell libraries were then created according to 10X Genomics specifications (Chromium Next GEM Single Cell 5ʹ v2.1-Dual Index Libraries). Samples were multiplexed and sequenced in pairs on an Illumina Novaseq X (Illumina, San Diego, CA). The sequenced data were processed into expression matrices with the Cell Ranger Single-cell software 9.0.0 obtained at the following website (https://www.10xgenomics.com/support/software/cell-ranger/latest/release-notes/cr-release-notes). FASTQ files were then obtained from the base-call files from Novaseq X sequencer and subsequently aligned to the rat genome NCBI Rnor6.0, with a read length of 26 bp for cell barcode and unique molecule identifier (UMI) (read 1), 8 bp i7 index read (sample barcode), and 98 bp for actual RNA read (read 2). Each rat sample yielded approximately 300 M reads.

For targeted analysis, Anndata^56^ and Scanpy^57^ were used to load and preprocess h5ad files for each sample for import into Seurat. Scrublet^58^ was used to identify and remove putative doublets before AnnData objects were concatenated by week. Barcodes, features, matrix files, and metadata were extracted into a folder for import into R with Seurat’s Read10X function^59^. 14-week and 26-week Seurat objects were created with CreateSeuratObject. Mitochondrial gene percentage was calculated for each cell in each Seurat object before removing cells containing greater than 5% mitochondrial reads. Mitochondrial genes and genes with fewer than 10 reads across all cells in each object were also removed. Lastly, any cells with fewer than 500 reads were removed. The 14- and 26-week Seurat objects were merged by timepoint and layers joined. Subsets of cardiomyocyte compartments were selected based on cell_type_leiden0.6 column. Samples were split by week and log-normalized with NormalizeData. FindVariableFeatures was used to identify the top 2000 variable features per week and the features were centered and scaled with ScaleData. Principle Component Analysis (RunPCA function) was run on the split objects with default parameters before layers were integrated back together (IntegrateLayers function) with method=RPCAIntegration. A shared nearest-neighbor graph was created with FindNeighbors (using 15 RPCA dimensions) and clustered with FindClusters (with a resolution of 0.5). Clusters were assigned cell types based on known marker genes. To create the heatmaps in Figure 2, the integrated Seurat object was subset into separate Endothelial, Myocyte, Fibroblast, and Pericyte objects. Seurat’s FindMarkers function was used to identify differentially expressed genes in ZSF1 Obese cells in each cell type, with a Bonferroni-corrected p-value cutoff of 0.05. Results were intersected with a list of genes involved in NO signaling and regulation of protein *s*-nitrosylation. Then the average Log_2_Fold-Change (FC) was plotted with ggplot2’s geom_tile function^60^.

### Gene Expression

mRNA expression profiles were assayed from heart, liver, and kidney tissue. Following RNA extraction, quantitative real time PCR was performed to probe for the mRNA expression. For eNOS (#4453320, ThermoFisher, Waltham, MA), iNOS (#4331382, ThermoFisher, Waltham, MA), nNOS (#4351372, ThermoFisher, Waltham, MA), and housekeeping 18S ribosomal RNA (#4333760, ThermoFisher, Waltham, MA), commercially available probes were utilized. Primer sequence for other tested genes (HBb, Trx2, and GSNOR) were listed below. 2^^delta-deltaCT^ values corrected to 18S rRNA were used for data analysis.

HBb: F-ACCTTGGCAGCCTCAGTG; R-GTGAATTCTTTGCCCAGGTGG.
Trx2: F-TCACACAGACCTTGCCATTGAG; R-CCTGCTTGTCAGCCAATTAG.
GSNOR: F-ACAGTGTGGAGAATGCAAG; R-GCTGGTTCCCATGAAGTG.

### Western Blot Analysis

Tissue were lyophilized with mortar and pestle and then homogenized in lysis Buffer 150mM NaCl (Calbiochem, 7760), 1% NP-40 (Sigma-Aldrich, 74385), 0,5% Na-deoxycholate (AppliChem, A1531,0025), 0,1% SDS (PanReac AppliChem, A2572), 50mM Tris-HCL, pH=7,4 (Sigma-Aldrich, T1503), 2mM EDTA (Merck, 4005) supplemented with a cocktail of protease (PI, Roche, 5892970001) and phosphatase inhibitors (PhoI, Roche, 4906837001). Lysates were centrifuged (13.000 rpm, 15min, 4-degree Celsius) and the protein concentration in the supernatants was quantified using the DC protein assay (BIO-RAD, 5000116). Concentration was normalized before Western blot analysis. Samples were separated on 10% or 12% SDS–PAGE and transferred to a nitrocellulose membrane (Macherey-Nagel; Düren, Germany), after Laemmli buffer containing 4% SDS, 10% β-mercaptoethanol (Sigma-Aldrich, M6250), 20% glycerol (Melford, GI345), 0,004% blue bromophenol (AppliChem, A2331,0025) and 0,125M Tris-HCL, was added. The membranes were blocked (5% milk (PanReac AppliChem, A0830)) and probed with the following antibodies: anti-β-Tubulin (Abcam, ab15568), anti-GAPDH (Proteintech, 10494-1-AP or Cell Signaling 2118S), anti-eNOS (Cell signaling, 32027s), anti-peNOSs1177 (Cell signaling, 9571), anti-HBb (abcam, ab214049), anti-Trx2 (abcam, ab185544), anti-GSNOR (Proteintech, 11051-1-AP). Immunoblots were next processed with anti-rabbit secondary antibody (Merck, AP132P) and visualized using the Western HRP substrate. In one subset of western blot experiments, membranes were incubated with No-Stain™ Protein Labeling Reagent (Invitrogen™, Waltham, MA) to visualize total protein as loading control. Quantification and normalization of Western blots was performed using ImageJ software (NIH Image, National Institutes of Health, USA). In the subset that utilized total protein as loading control, target proteins were normalized to the signal from entire lane; however, only a portion of the membrane around the same molecular weight as the target proteins were shown due to limited space allowed.

### Thioredoxin Activity Measurements

Trx activity in tissue were measured using a commercially available fluorometric activity assay kit (#500228, Cayman Chemical, Ann Harbor, MI) following the instructions by the manufacturer in combination with a Synergy LX multi-mode microplate reader (Agilent, Santa Clara, CA).

### GSNOR Activity Measurements

GSNOR activity in tissue were measured as previously described^36, 61^. Briefly, tissue samples were homogenized in a buffer of 50 mmol/l Tris-HCL (pH 8.0), 150 mM NaCl, 1 mM EDTA, 0.1% Triton and 1:100 protease inhibitor cocktail (Sigma). The samples were then centrifuged at 10,000 RCF for 10 minutes at 4 degrees Celsius and brought to a concentration of 0.1 mg/ml in reaction buffer of 20 mmol/l Tris-HCl, 0.5 mmol/l EDTA. Samples were then added 75 µM NADH with or without 100 µM of GSNO. Samples were measured using fluorescence at 350 nm excitation / 460 nm emission. Every minute for one hour where the rate is determined by NADH consumption with and without the GSNO substrate.

### *Ex Vivo* Aortic Ring Vascular Reactivity

Thoracic aortas of the rats were removed for vascular reactivity testing at sacrifice as previously described^53, 54^. The rings were first equilibrated in Krebs-Henseleit solution and provided a tension of 0.5 grams for 60 minutes to reach equilibration. They were then pretreated with phenylephrine for maximal constriction and then followed with challenges of titrated acetylcholine and subsequently sodium nitroprusside and measured relaxation as compared to phenylephrine maximal contraction.

### Statistical Analysis

All data in this study are presented as mean ± SEM. All statistical analysis were performed in a blinded manner using the Prism 10 software (GraphPad, San Diego, California). Normal distribution was tested by the D’Agostino & Pearson test. Differences in data among 2 groups were compared using an unpaired student t-test, or a Mann Whitney test if the data did not follow a normal distribution or violated the assumption of equal variance. For multiple group comparison over time, 2-way ANOVA was utilized. For dataset involving multiple factors such as timepoints or concentrations, 2-way ANOVA analysis followed by a Bonferroni multiple comparison test was utilized. A p value of <0.05 was considered statistically significant. The presented data may have different numbers of animals per group as only a subset of the mice from each group were used for certain experiments. Additionally, exclusion of animals was carried out due to complications such as procedural failure in invasive hemodynamics measurement, limited amount of sample collected (i.e plasma volume), and lack of participation in treadmill running. Prior to conducting statistical analysis, an outlier test was performed using “ROUT” method developed by Prism 10 to identify and remove any outliers in the data set.

## RESULTS

### Increased RsNO and 3-NT in plasma samples of HFpEF patients

To evaluate NO bioavailability as well as protein *s-*nitrosylation, and nitrosative stress we measured plasma nitrite, RsNO, and 3-NT in HFpEF patients and controls. Nitrite is a well-established biomarker of biologically active NO, while RsNO and 3-NT are biomarkers for protein *s*-nitrosylation and nitrosative stress, respectively^62–64^. We observed that plasma nitrite levels remained unchanged in HFpEF patients (***Figure 1A***). Despite a lack of evidence for increase NO bioavailability as circulating nitrite levels remained unaltered, circulating RsNO levels were significantly elevated in HFpEF patients indicating increased protein nitrosylation (***Figure 1B***). Importantly, 3-NT was increased by greater than 2-fold in HFpEF patients as compared to control subjects (***Figure 1C***). These data provide strong evidence for pathological protein *s-*nitrosylation and nitrosative stress in HFpEF patients despite no alterations in NO bioavailability.

**Figure 1.**
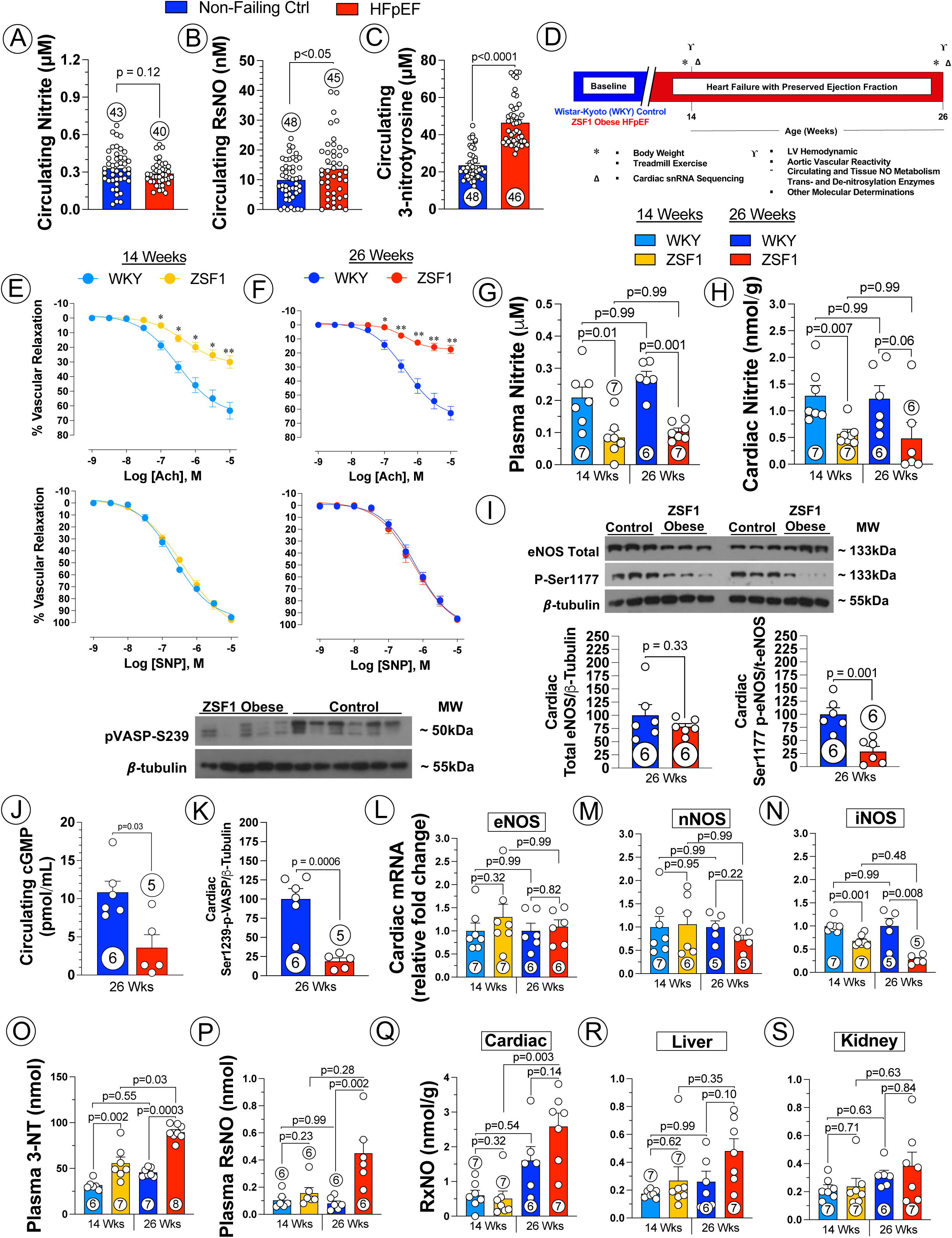
Dysregulated NO Metabolites in Patients with HFpEF and ZSF1 obese Rat Model of HFpEF. Nitrite ***(A)***, *s*-nitrosothiols (RsNO) ***(B)***, and 3-nitrotyrosine ***(C)*** levels in plasma samples collected from control vs. HFpEF patients. Study timeline of the 12-week study in ZSF1 Ob rat model of HFpEF (***D***). Vasorelaxation response curves to endothelium-dependent vasodilator acetylcholine (ACh) or endothelium-independent vasodilator sodium nitroprusside (SNP) in aorta isolated from WKY control vs. ZSF1 obese rats at 14 *(**E**)* and 26 *(**F**)* weeks of age. Nitrite levels in plasma *(**G**)* and cardiac tissue *(**H**)* from ZSF1 Ob vs. WKY rats at 14 or 26 weeks of age. Representative Western blot as well as protein expression quantification for total eNOS and P-Ser1177 eNOS *(**I**)* in cardiac tissues of 26-week-old WKY vs. ZSF1 obese rats. Circulating cGMP (***J***) level in 26-week-old WKY vs. ZSF1 Ob rats. Cardiac phosphorylated VASP (***K***) in cardiac tissues of 26-week-old WKY vs. ZSF1 obese rats. mRNA levels of eNOS *(**L**)*, nNOS *(**M**)*, and iNOS *(**N**)* in the cardiac tissue from WKY control vs. ZSF1 obese rats. Circulating 3-NT (***O***) and RsNO (***P***) levels in 14-week-old and 26-week-old ZSF1 Ob rats vs. WKY controls. Cardiac (***Q***), hepatic (***R***), and renal (***S***) RxNO levels in 14-week-old and 26-week-old ZSF1 Ob rats vs. WKY controls. In Figure 1A ***to 1E*** and ***1H to 1J***, data were analyzed with student unpaired 2-tailed t test. Data were presented as mean ± SEM. Circled number inside bars indicate sample size.

**Figure 2.**
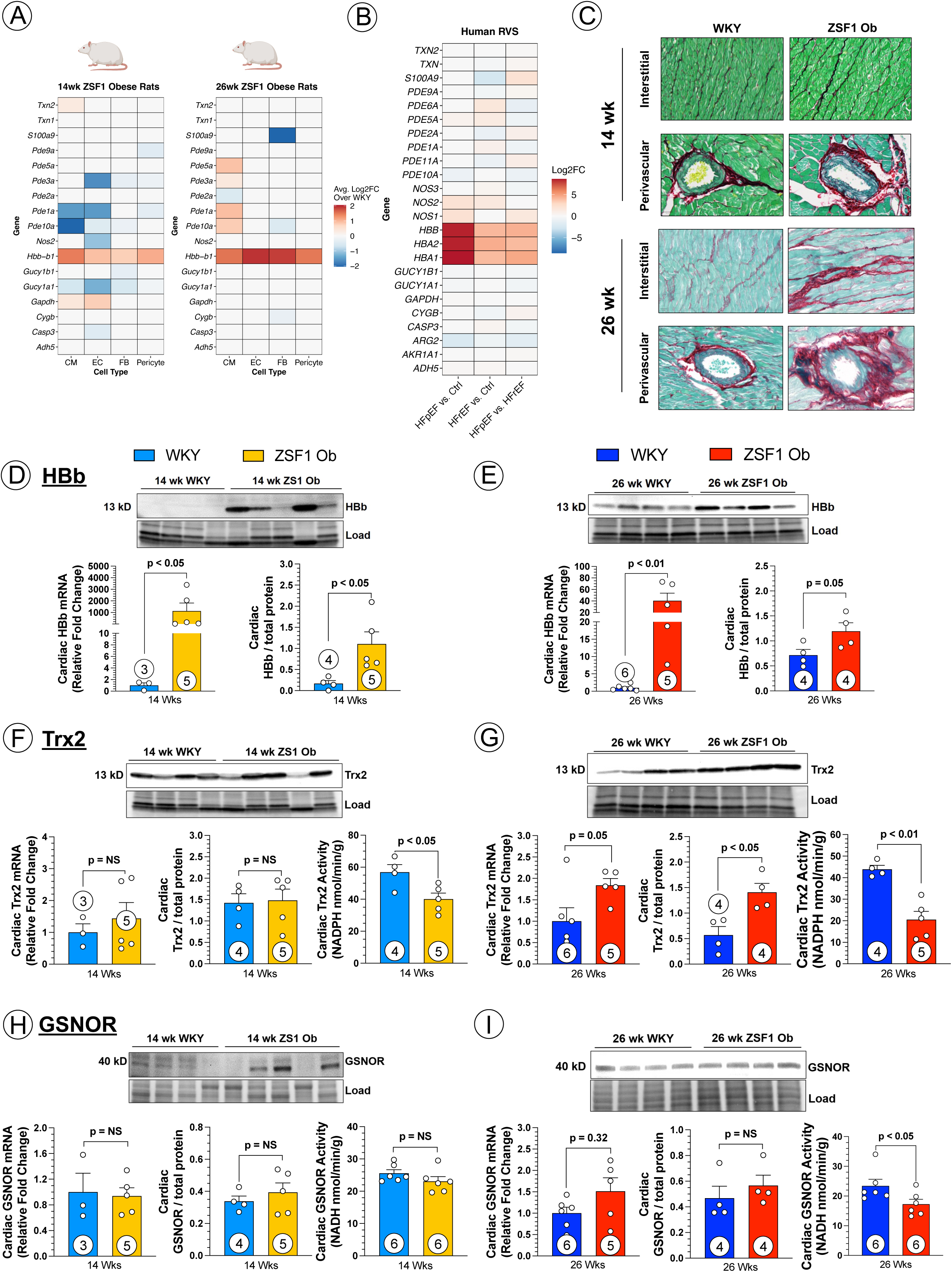
Imbalance of Trans-nitrosylase (HBb expression) and De-nitrosylase in ZSF1 Ob Heart. Heat map of average Log_2_Fold-Change (FC) for NO synthase, selected downstream effectors, and common trans- and de-nitrosylases acquired via targeted analysis of single nuclei RNA sequencing of cardiac tissues from 14-week-old and 26-week-old WKY controls vs. ZSF1 obese rats (***A***). Heat map of Log_2_FC for NO synthase, selected downstream effectors, and common trans- and de-nitrosylases acquired via targeted analysis of bulk RNA sequencing of cardiac biopsies collected from HFpEF, HFrEF, and control patients (***B***). Representative microphotography of cardiac fibrosis in interstitial area and perivascular area from ZSF1 Ob vs. WKY hearts at 14 or 26 weeks of age (***C***). mRNA, representative western blot image and quantified protein expression of HBb in cardiac tissues collected from 14-week-old *(**D**)* and 26-week-old *(**E**)* WKY controls vs. ZSF1 obese rats. mRNA, representative western blot image and quantified protein expression, and enzymatic activity of Trx2 in cardiac tissues collected from 14-week-old *(**F**)* and 26-week-old *(**G**)* WKY controls vs. ZSF1 obese rats. mRNA, representative western blot image and quantified protein expression, and enzymatic activity of GSNOR in cardiac tissues collected from 14-week-old *(**H**)* and 26-week-old *(**I**)* WKY controls vs. ZSF1 obese rats. Data in Figure 2A was analyzed with 2-way ANOVA followed by Bonferroni post-hoc analysis; data in ***2B*** were analyzed with 1-way ANOVA; in ***Figure 2D - I***, data were analyzed with Mann Whitney test and presented as mean ± SEM. Circled number inside bars indicate sample size.

### ZSF1 obese rats develop severe cardiometabolic HFpEF

Male WKY control and ZSF1 obese rats were monitored for a period of 12 weeks starting at the 14 weeks of age to evaluate the progression of HFpEF in ZSF1 obese rats. At 14 and 26 weeks of age, body weight, echocardiographic parameters, treadmill exercise performance were measured (***Supplemental Figure 1***). A subset of rats was assessed for LV and systemic hemodynamics and aortic vascular reactivity, followed by sacrifice for other molecular determinations. An illustrative protocol is depicted in ***Figure 1D***.

Throughout the 12-week study, ZSF1 obese rats displayed markedly higher body weight as compared to WKY control rats (***Supplemental Figure 1A***), which was accompanied by significantly impaired glucose handling ability and reduced insulin sensitivity (data not shown), as depicted previously^41, 43^. Invasive hemodynamic measurements revealed that ZSF1 obese rats exhibited severe hypertension with significantly elevated systolic and diastolic blood pressures throughout the study (***Supplemental Figure 1B***). Hypertension in the ZSF1 rat was accompanied by severe cardiac hypertrophy as measured in heart weight/tibia length in ZSF1 obese rats (***Supplemental Figure 1C***). Importantly, we observed significant elevations in LV end-diastolic pressure (***Supplemental Figure 1D***) and Tau (***Supplemental Figure 1E***) in ZSF1 obese rats, indicating severe cardiac diastolic dysfunction, one of the hallmarks of HFpEF. Severe exercise intolerance was also observed in ZSF1 obese rats as measured in treadmill running distance during exercise (***Supplemental Figure 1F***).

### ZSF1 obese rat exhibited dysregulated eNOS activity and signaling and reduced NO bioavailability

Endothelial function and eNOS - derived NO signaling have long been considered crucial for cardiovascular hemostasis, and reduced NO signaling has been linked to diastolic dysfuction^65^. To assess the extent of dysregulation in NO signaling, we first measured the eNOS-dependent vascular function in isolated, thoracic aortic ring segments. Throughout the 12-week study, thoracic aortic rings from ZSF1 Ob rats displayed significantly blunted responses to the endothelial-dependent vasodilator, acetylcholine (ACh) as compared to the WKY control group. In contrast, vascular reactivity to the direct nitro-vasodilator, sodium nitroprusside (SNP) remained unchanged (***Figures 1E - F***). Endothelial dysfunction and attenuated NO bioavailability were further corroborated by significantly reduced nitrite levels in the circulation and myocardial tissue (***Figures 1G - H***). Further, we observed no difference between ZSF1 rats vs. WKY control rats in total cardiac eNOS expression but a significant reduction in phosphorylation on the serine 1177 residues of cardiac eNOS, an eNOS activation site, at 26 weeks of age, confirming the disruption in eNOS activity (***Figure 1I***). Downstream effectors of eNOS activation, circulating cGMP and PKG-mediated phosphorylation of vasodilator-stimulated phosphoprotein (pVASP) were also significantly reduced in ZSF1 Ob rats (***Figures 1J - K***). Taken together, these results indicate profound endothelial dysfunction and eNOS dysregulation accompanied by significant reductions in NO bioavailability. Additionally, we measured the mRNA levels of eNOS, iNOS and nNOS isoforms. There was no difference observed in eNOS and nNOS mRNA levels throughout the study (***Figures 1L - M***). Interestingly, iNOS mRNA levels were reduced in ZSF1 rats throughout the study while its protein expression was reduced at 14 weeks of age, but remained unchanged at 26 weeks of age (***Figure 1N, Supplemental Figure 2A - B***).

### ZSF1 obese rats displayed systemic elevation in nitrosative stress and protein *s*-nitrosylation

To evaluate the extent of nitrosative stress in ZSF1 obese rats, we measured 3-NT and RsNO in the plasma, as well as RxNO in the heart. Starting at 14 weeks of age ZSF1 Ob HFpEF rats exhibited age-dependent increases in both the 3-NT and RsNO levels compared to WKY control rats (***Figures 1O - P***). In contrast, there was no difference in cardiac RxNO levels at the early stage of HFpEF development (ie., 14 weeks of age); however, at 26 weeks of age, ZSF1 obese rats displayed significantly elevated levels of RxNO in the heart, indicating a significant increase in myocardial protein *s-*nitrosylation (***Figure 1Q***). To better understand the extend of such elevated nitrosative stress and protein *s*-nitrosylation, we next measured RxNO levels in additional organs including the liver and the kidney, both of which are commonly linked to HFpEF comorbidities and complications. Similar to the heart, hepatic and renal RxNO levels were elevated in an age-dependent manner in ZSF1 Obese rat model of HFpEF (***Figures 1R - S***).

### Imbalance of trans- and de-nitrosylases underpinned the elevated cardiac nitrosative stress in ZSF1 obese rat

Next, we performed snRNA Seq to acquire molecular insights into the development of elevated cardiac nitrosative stress despite the reduction in NO bioavailability and signaling. Targeted analysis revealed changes in the expression of a variety of genes related to NO generation and signaling in 4 cardiac resident cell types including cardiomyocytes, endothelial cells, fibroblasts, and pericytes. No change was observed in eNOS or nNOS expression and iNOS was significantly down-regulated in myocardial endothelial cells in early HFpEF (i.e. 14 weeks of age) (***Figure 2A***). Notably, expression of hemoglobin subunit beta (HBb), a well characterized transnitrosylase^66, 67^, was markedly elevated in all 4 resident cardiac cell types at both the early and late stages of HFpEF (***Figure 2A***). In contrast, expression of common de-nitrosylases such as thioredoxins, GSNOR, AKR1a1 remained relatively stable as the only change observed was a mildly increased Trx2 expression in cardiomyocytes in 14-week-old ZSF1 Ob rats (***Figure 2A***). Importantly, we confirmed that our observations made in ZSF1 Ob rat model of cardiometabolic HFpEF were true in human HFpEF patients^1^. Targeted analysis of the transcriptome of cardiac biopsies from HFpEF patients^1^ demonstrated a marked upregulation of HBb when compared to either control or HFrEF (***Figure 2B***). Taken together, our data reveal a significant imbalance in trans- and de-nitrosylase expression, particularly in HBb expression, suggesting that this imbalance may be the primary culprit responsible for elevated cardiac protein *s*-nitrosylation and subsequent nitrosative stress.

To further examine this hypothesis, we performed real time PCR, western blot, and activity assay experiments to measure HBb, Trx2, and GSNOR in cardiac tissue homogenates. Associated with substantial formation of cardiac interstitial and perivascular fibrosis (***Figures 2C***), and consistent with the snRNA Seq analysis, striking increases in cardiac HBb mRNA and protein levels were observed in ZSF1 Ob rats at both 14 and 26 weeks of age (***Figures 2D - E***). mRNA and protein levels of Trx2 remained unaltered in ZSF1 Ob hearts at 14 weeks of age but significantly increased at 26 weeks of age; however, Trx enzyme activity was downregulated in ZSF1 Ob hearts at both 14 and 26 weeks of age (***Figures 2F - G***). Similarly, despite no change mRNA and protein levels, cardiac GSNOR activity was significantly reduced in the ZSF1 Ob heart at 26 weeks of age as compared to WKY control (***Figures 2H - I***).

### Imbalance of trans- and de-nitrosylases contributed to elevated RxNO in peripheral organs in ZSF1 obese rat

Given that HFpEF comorbidities (i.e., hypertension, obesity, diabetes) impact multiple organ systems and the pathophysiology of HFpEF involves organs beyond the heart and circulation, we evaluated the involvement of trans- and de-nitrosylases (i.e., HBb, Trx2, and GSNOR) in the liver and kidney in ZSF1 obese rats. Associated with significant hepatic steatosis found in ZSF1 Ob liver (***Figures 3A - B***), we observed markedly elevated hepatic HBb mRNA and protein levels throughout the 12-week study (***Figures 3C - D***). Hepatic Trx2 mRNA level was down-regulated at 14 weeks of age, but protein expression and enzymatic activity was unaffected. While at 26 weeks of age, hepatic Trx2 protein increased but the enzymatic activity was reduced in ZSF1 Ob rats (***Figures 3E - F***). Conversely, GSNOR mRNA, protein, and enzymatic activity were all significantly reduced in ZSF1 Ob hepatic tissue at both 14 and 26 weeks of age (***Figures 3G - H***).

**Figure 3.**
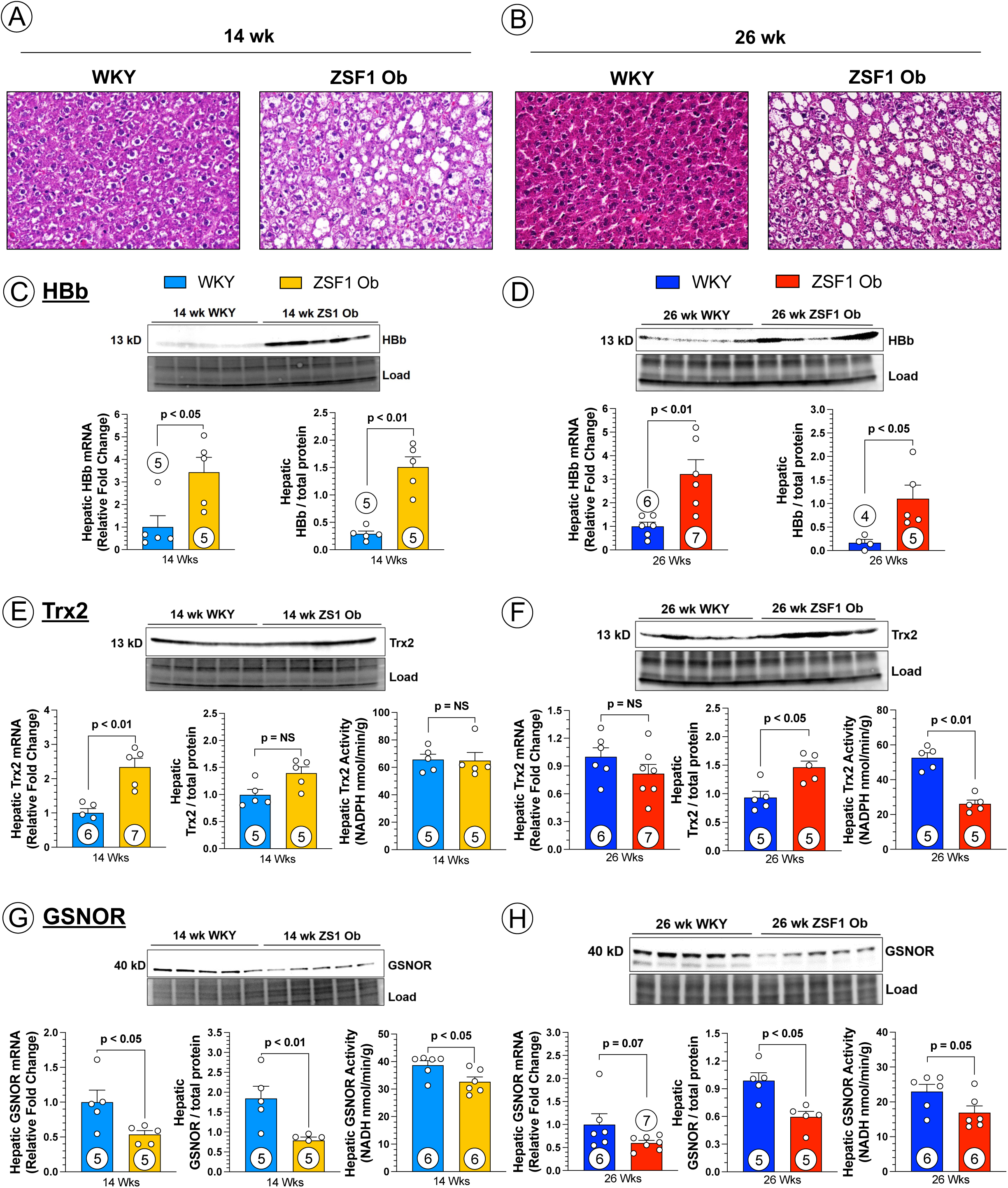
Imbalance of Trans-nitrosylase (HBb expression) and De-nitrosylase in ZSF1 Ob Liver. Representative microphotography of hepatic lipid deposition from ZSF1 Ob vs. WKY livers at 14 (***A***) or 26 weeks (***B***) of age. mRNA, representative western blot image and quantified protein expression of HBb in hepatic tissues collected from 14-week-old *(**C**)* and 26-week-old *(**D**)* WKY controls vs. ZSF1 obese rats. mRNA, representative western blot image and quantified protein expression, and enzymatic activity of Trx2 in hepatic tissues collected from 14-week-old *(**E**)* and 26-week-old *(**F**)* WKY controls vs. ZSF1 obese rats. mRNA, representative western blot image and quantified protein expression, and enzymatic activity of GSNOR in hepatic tissues collected from 14-week-old *(**G**)* and 26-week-old *(**H**)* WKY controls vs. ZSF1 obese rats. Data were analyzed with Mann Whitney test and presented as mean ± SEM. Circled number inside bars indicate sample size.

In association with the age-dependent formation of tubulointerstitial and perivascular fibrosis in ZSF1 Ob kidneys (***Figures 4A - B***), a profound increase in HBb mRNA and protein levels was observed in ZSF1 Ob kidneys at both 14 and 26 weeks of age, consistent with those demonstrated in the cardiac and hepatic tissue (***Figures 4C - D***). Renal Trx2 mRNA was reduced in ZSF1 Ob rats at 26 but not 14 weeks of age; however, renal Trx2 protein was increased significantly at 26-but not 14-week-old ZSF1 Ob rats. In contrast, renal Trx2 enzymatic activity was significantly reduced in both 14- and 26-week-old ZSF1 Ob HFpEF rats (***Figures 4E - F***). Conversely, renal GSNOR mRNA and protein levels were reduced at both 14 and 26 (significantly) weeks of age, while GSNOR activity were significantly reduced only in 26-week-old ZSF1 Ob rats (***Figures 4G - H***).

**Figure 4.**
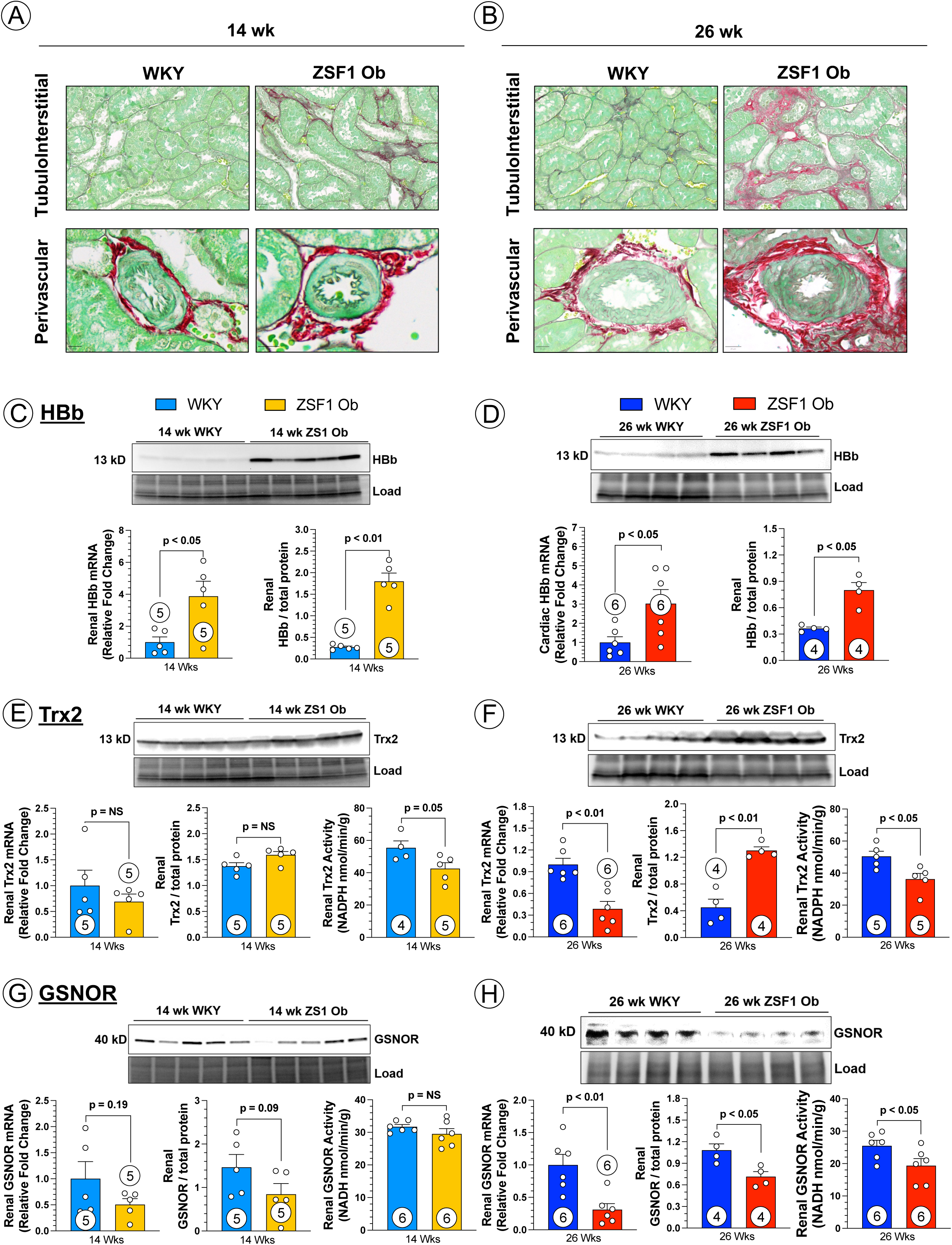
Imbalance of Trans-nitrosylase (HBb expression) and De-nitrosylase in ZSF1 Ob Kidney. Representative microphotography of renal fibrosis in tubulointerstitial area and perivascular area from ZSF1 Ob vs. WKY kidneys at 14 (***A***) or 26 weeks (***B***) of age. mRNA, representative western blot image and quantified protein expression of HBb in renal tissues collected from 14-week-old *(**C**)* and 26-week-old *(**D**)* WKY controls vs. ZSF1 obese rats. mRNA, representative western blot image and quantified protein expression, and enzymatic activity of Trx2 in renal tissues collected from 14-week-old *(**E**)* and 26-week-old *(**F**)* WKY controls vs. ZSF1 obese rats. mRNA, representative western blot image and quantified protein expression, and enzymatic activity of GSNOR in renal tissues collected from 14-week-old *(**G**)* and 26-week-old *(**H**)* WKY controls vs. ZSF1 obese rats. Data were analyzed with Mann Whitney test and presented as mean ± SEM. Circled number inside bars indicate sample size.

### Alterations in trans- and de-nitrosylases lead to dysregulated NO and RxNO dynamic in the “2-hit” murine HFpEF model

To further examine the dysregulated dynamic of NO and RxNO, and the role of trans- and de-nitrosylases in the setting of HFpEF, we performed additional experiments in another well-established rodent model of cardiometabolic HFpEF, the “two-hit” murine model induced by L-NAME and high fat diet (HFD) treatment (***Supplemental Figure 3A***)^22, 46, 47, 68^. Mice treated with L-NAME and HFD displayed significantly lowered plasma and cardiac tissue levels of nitrite, indicating a pathological reduction in NO bioavailability (***Supplemental Figures 3B - C***), similar to observations in the ZSF1 obese rat. Interestingly, plasma 3-NT and RsNO, were significantly elevated in the “2-Hit” mouse model of HFpEF (***Supplemental Figures 3D - E***). Following 10 weeks of “Two-hit”, RxNO levels were also markedly elevated in the heart and various peripheral organs including the liver, kidney and lung (***Supplemental Figure 5F***), highlighting the paradoxical systemic imbalance of NO bioavailability and excessive nitrosative stress. Correspondingly, HBb protein expression was upregulated in the heart, liver, and kidney of “Two-hit” HFpEF mice (***Supplemental Figures 3G - J***). Conversely, GSNOR protein expression was not altered in the heart and kidney, while significantly increased in the liver of the “two-hit” HFpEF mice (***Supplemental Figures S3K - N***). Despite unchanged protein levels, enzymatic activity of GSNOR was reduced in the heart, liver, kidney, and lung of the “2-Hit” HFpEF mice when compared to control (***Supplemental Figure 5O***).

## DISCUSSION

HFpEF is a heterogenous, multi-organ disease associated with high morbidity and mortality^2, 5, 69^. Recently the prevalence of HFpEF has increased significantly and HFpEF has now eclipsed HFrEF in terms of new HF diagnoses. There are multiple classes of effective therapeutic treatments accessible for patients with HFrEF, yet these treatments are largely ineffective for treating HFpEF ^70^. Thus, despite similar symptoms, there are likely very distinct underlying cellular and molecular pathological mechanisms that distinguish HFrEF from HFpEF^70^. Elucidation of these divergent mechanism is necessary to identify effective therapeutic targets to aid in the development of effective therapies for the treatment of HFpEF.

Given the well-established role of NO signaling in the regulation of cardiovascular health and the numerous reports demonstrating reduced NO bioavailability in CV diseases, NO-based therapies have been tested in HFpEF ^18–21^. However, this therapeutic approach has failed to result in meaningful clinical efficacy in HFpEF patients. Clinical studies utilizing nitrite, nitrate, or organic nitrates to replenish NO bioavailability, or sGC stimulators and PED5 inhibitors to amplify NO-mediated cellular signaling have produced neutral results, despite the plethora of evidence suggesting that depleted NO bioavailability as a hallmark feature of HFpEF^13^. However, it is possible that increased superoxide radical anions that occurs in HFpEF conjugates with the supplemental NO to form peroxynitrite, significantly suppressing the therapeutic action of NO-based therapy. Indeed, we have previously reported that NO-based therapy (i.e, sodium nitrite), when combined with a powerful antioxidant and peroxynitrite scavenger, hydralazine, was highly effective in reducing the severity of HFpEF in the “2-Hit” mouse model^47^. Such approach warrants further exploration in both basic and clinical research settings.

Additionally, recent preclinical studies have suggested that excessive nitrosative stress and accumulation of nitroso - species (RxNO) such as *s*-nitrosylated proteins play a causative role in the pathogenesis and progression of HFpEF^22, 23^. Schiateralla et al. described an increase in iNOS levels coupled with observation of increased cardiac protein *s*-nitrosylation (in particular cardiac IRE1α), which in turn, participated in the pathogenesis of HFpEF in the “two hit” mouse model of HFpEF^22^. Moreover, utilizing a different animal model, the salty drinking water/unilateral nephrectomy/aldosterone (SAUNA) rat model of HFpEF, Yoon et al. demonstrated that upregulation of nNOS, instead of iNOS, was responsible for increased *s*-nitrosylation of histone deacetylase 2, which in turn, induced cardiac diastolic dysfunction^23^. These landmark studies established the possible link between nitrosative stress as well as pathological accumulation of RxNO in HFpEF, providing an alternative explanation on the neutral results produced by NO-based therapies for the treatment of HFpEF. In the present study, we sought to confirm the previous findings of elevated NO production by NOS and excessive nitrosative stress in the setting of HFpEF. Our initial studies in HFpEF patients revealed for the first time that plasma RsNO and 3-NT are significantly elevated in HFpEF patients as compared to control, which is indicative of excessive nitrosative stress in agreement with previous reports^22, 23^; however, we did not observe any changes in circulating nitrite, a well-established biomarker and sink of systemic NO bioavailability. These findings inform the potential use of RsNO and 3-NT as circulating biomarkers for the diagnosis of HFpEF following comprehensive clinical testing in large, diverse patient populations. Furthermore, we evaluated nitrite and RxNO bioavailability in two well established rodent models of cardiometabolic HFpEF, the ZSF1 obese rat model and the “two-hit” mouse model ^22, 40, 42^. In concert with our observations in clinical samples, ZSF1 obese rats displayed significantly increased levels of *s*-nitrosylated proteins in both the circulation and other organs during the progression of HFpEF. Increased RxNO levels observed in the heart as well as peripheral organs such as the liver, kidney, and the lung (mouse), suggest that nitrosative stress is a systemic pathological component of HFpEF. However, in both ZSF1 Ob rat and “two-hit” mouse model, we observed profound reductions in nitrite levels in the circulation, cardiac and other tissue (data not shown). Together with the observations of impaired endothelial-dependent vasodilation in response to acetylcholine treatment, reduced phosphorylated eNOS (serine 1177), decreased circulating cGMP, and lower cardiac pVASP, these findings conclusively demonstrate that NO bioavailability and signaling are severely impaired in these models of HFpEF. Taken together, these data identified a paradoxical dysregulation of NO - RxNO dynamic characterized by impaired NO production and signaling coupled with excessive circulating and tissue protein *s*-nitrosylation. Notably, such phenomenon is contradictory to what was observed in the context of other cardiovascular diseases including myocardial infarction and HFrEF, in which NO availability and signaling is similarly reduced while overall protein *s*-nitrosylation is largely insufficient^15, 36, 71–77^. These novel findings suggest that increased production of NO may not be the cause of nitrosative stress in HFpEF, but that an alternative cellular process is responsible for the excess accumulation of RxNO/nitrosative stress.

Indeed, the levels of these nitroso - species are also delicately regulated by the trans-nitrosylases, enzymes that transfer NO onto the cysteine residues of proteins, and by the denitrosylases, enzymes that remove the NO moiety from the cysteine residues^27, 28, 33, 78, 79^. Similar to other post-translational protein modifications such as phosphorylation and sulfhydration, hyper- or hypo-*s*-nitrosylation of specific proteins resulted in altered function, which could lead to a wide range of effects that impact disease progression^27, 32, 33^. Our findings indicate that excessive protein *s*-nitrosylation likely contributes to HFpEF disease progression, given the elevation Rx(s)NO and 3-NT levels in multiple organs of animal models and in HFpEF patients. However, the roles of trans- and de-nitrosylases, and their intricate interplay with bioactive NO has not been investigated in the context of HFpEF. Utilizing snRNA seq, we unexpectedly determined that hemoglobin (subunit beta), a trans-nitrosylase enzyme that normally expresses mainly in erythrocytes, was shown to be markedly upregulated in all four major resident cardiac cell types, cardiomyocytes, endothelial cells, fibroblasts, and pericytes. Corroborating the snRNA seq analysis, we also observed striking upregulation of HBb in mRNA and protein levels in cardiac, hepatic, and renal tissues. To determine whether this observation was true in human HFpEF patients, we examined the data from a landmark study by Hahn et al. investigating the myocardial gene expression signature in human HFpEF and found that cardiac biopsies from HFpEF patients displayed an over 256-fold increase in HBb when compared to control subjects, and an over 64-fold increase when compared to HFrEF patients^1^. It is well-established that hemoglobin is not normally expressed in appreciable levels in the myocardium and other peripheral organs^66, 71, 80–83^. We are aware of only a few studies describing elevated hemoglobin expression in non-erythrocytes such as cardiac and hepatic tissues triggered by a pro-oxidative and inflammatory cellular environment^84–86^. Our findings provide novel insights into the potential deleterious roles of hemoglobin in cardiometabolic HFpEF. Specifically, we proposed that elevated HBb across multiple organs might contribute to HFpEF pathophysiology by two distinct mechanisms. First, hemoglobin acts as an NO scavenger by readily binding to it with high affinity, thereby eliminating NO bioavailability and impairing NO signaling^87–91^, which could explain the reduced nitrite levels in both rodent HFpEF models. Second, it is plausible that such profound upregulation of HBb substantially enhanced overall trans-nitrosylation processes in the heart, liver, and kidney, thereby contributing to the excessive RxNO observed in these key organs and driving the pathogenesis and progression of cardiometabolic HFpEF.

Other key factors responsible for the dynamic control of protein *s*-nitrosylation are de-nitrosylases such as Trx and GSNOR, which are among the most ubiquitously expressed and well-characterized de-nitrosylases and responsible for de-nitrosylating a wide range of protein targets^26, 30, 78, 92, 93^. Trx, mainly known for its antioxidant property, has recently been shown to de-nitrosylate a wide range of protein targets in mammalian cells^92, 93^. snRNA seq analysis revealed that both Trx isoforms (Trx 1 in the cytosol and Trx 2 in the mitochondria) remained relatively stable, except a mild increase in Trx2 in ZSF1 Ob cardiomyocytes at 14 weeks of age (snRNA seq analysis showed Trx2 to be the predominant isoform in the heart). As a result, we further examined Trx2 mRNA, protein expression, and enzymatic activity levels in ZSF1 Ob rats. Interestingly, significantly reduced Trx enzymatic activity was observed in the cardiac, hepatic, and renal tissues, despite unaltered or increased protein expression. These data suggest that strategies enhancing Trx activity may be promising therapies in the context of cardiometabolic HFpEF, as such strategies likely results in dual beneficial actions in reducing both oxidative stress and nitrosative stress. Conversely, previous studies of GSNOR had focused on the therapeutic benefits of GSNOR inhibition. It has been shown that reduced GSNOR activity exerts beneficial effects in animal models of hypertension, atherosclerosis, and myocardial ischemia/reperfusion injury by promoting angiogenesis, inhibiting apoptosis, and reducing inflammation and oxidative stress^27, 30, 74–77^. However, recently emerged evidence suggests otherwise: deficiency in GSNOR resulted in excessive nitrosative stress, leading to cellular senescence, improper mitochondrial turnover, hepatic insulin resistance in diabetes and obesity, as well as aberrant placental *s*-nitrosylation and preeclampsia^34, 94, 95^. In the present study, we also observed reduced GSNOR protein expression and enzymatic activity in an age-dependent manner, which were associated with increased RxNO/nitrosative stress levels in multiple organs including the heart, liver, kidney, and lung (“two-hit” mouse model), informing possible therapeutic benefits of GSNOR activation instead of inhibition in the context of cardiometabolic HFpEF.

Taken together, we revealed for the first time that circulating markers of protein *s*-nitrosylation and nitrosative stress were increased in HFpEF patients, and that the equilibrium between endogenous NO production and signaling (i.e., NOS expression, activity, and downstream effectors) and enzymatic modulation of *s*-nitrosylation by trans- and de-nitrosylases pathways (e.g., HBb expresion, Trx2 and GSNOR expression and activity) is interrupted in the pathogenesis of cardiometabolic HFpEF in two well-characterized rodent models. These findings are the first evidence that dysregulated trans- or de-nitrosylases, instead of overproduction of NO by NOS, contribute to the pathological systemic accumulation of RxNO during the pathogenesis and progression of HFpEF, potentially leading to comorbidities in peripheral organs including the liver and kidney (***Figure 5***). Restoration of the balance between trans- and de-nitrosylase expression and activity represents a new therapeutic approach to combat the heightened nitrosative stress seen in cardiometabolic HFpEF patients.

**Figure 5.**
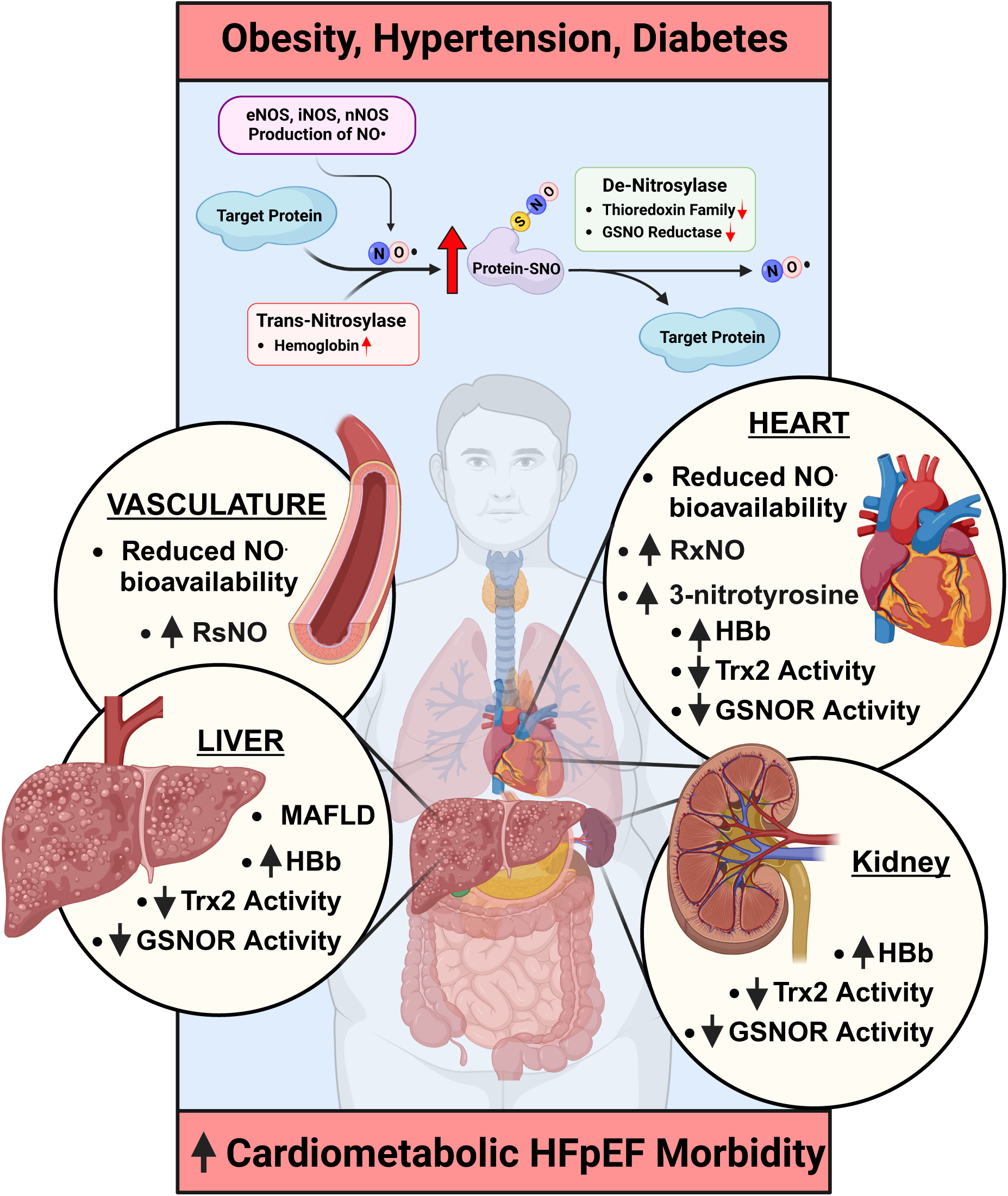
Working Hypothesis of Imbalance between NO bioavailability and Excessive Nitrosative Stress Caused by Dysregulated Trans- and De-Nitrosylases. The equilibrium between endogenous NO production/signaling (i.e., reduced eNOS activity and downstream effectors) and enzymatic modulation of *s*-nitrosylation by trans- and de-nitrosylases pathways (i.e., increased HBb expression, decreased Trx2 as well as GSNOR expression and activity) is interrupted in the pathogenesis of cardiometabolic HFpEF.

### Study Limitations

The current study utilized the ZSF1 obese rat and “two-Hit” mouse models of HFpEF. Although these rodent models have been well characterized, they feature metabolic dysfunction and hypertension which only represents a subset of all HFpEF patients. Such approach provided limited knowledge on the roles of trans- and de-nitrosylases including HBb, Trx2, and GSNOR in HFpEF with other HFpEF phenogroups. Development of novel HFpEF animal models that reflect diverse phenogroups is urgently needed to advance our understanding of HFpEF and to better design treatment strategies. Additionally, although we thoroughly examined the expression levels and enzymatic activity of these trans- or de-nitrosylases, future studies utilizing post-translational modification proteomic techniques are warranted to determine the target proteins that are subjected to hyper-nitrosylation. Finally, only male rats and mice were utilized in the current study due to the recent report suggesting female HFpEF patients are more likely to display lower body mass index as compared to their male counterpart^48^. Nevertheless, future studies aimed to investigate the roles of dynamic control of trans- and de-nitrosylation in female HFpEF are warranted.

## Nonstandard Abbreviations and Acronyms

3-NT: 3-nitrotyrosine
ACh: acetylcholine
EDP: end-diastolic pressure
EF: ejection fraction
eNOS: endothelial nitric oxide synthase
GSNOR: S-nitrosoglutahione reductase
HBb: Hemoglobin Subunit Beta
HFpEF: heart failure with preserved ejection fraction
HFrEF: heart failure with reduced ejection fraction
L-NAME: N(ω)-nitro-L-arginine methyl ester
LV: left ventricle
NO: nitric oxide
nNOS: neuronal nitric oxide synthase
RsNO: nitrosothiols
RxNO: total nitroso - species
SNP: sodium nitroprusside
Trx: Thioredoxin
WKY: wistar kyoto rat
ZSF1 Ob: zucker fatty and spontaneously hypertensive obese rat

---

**Supplemental Figure 1.**
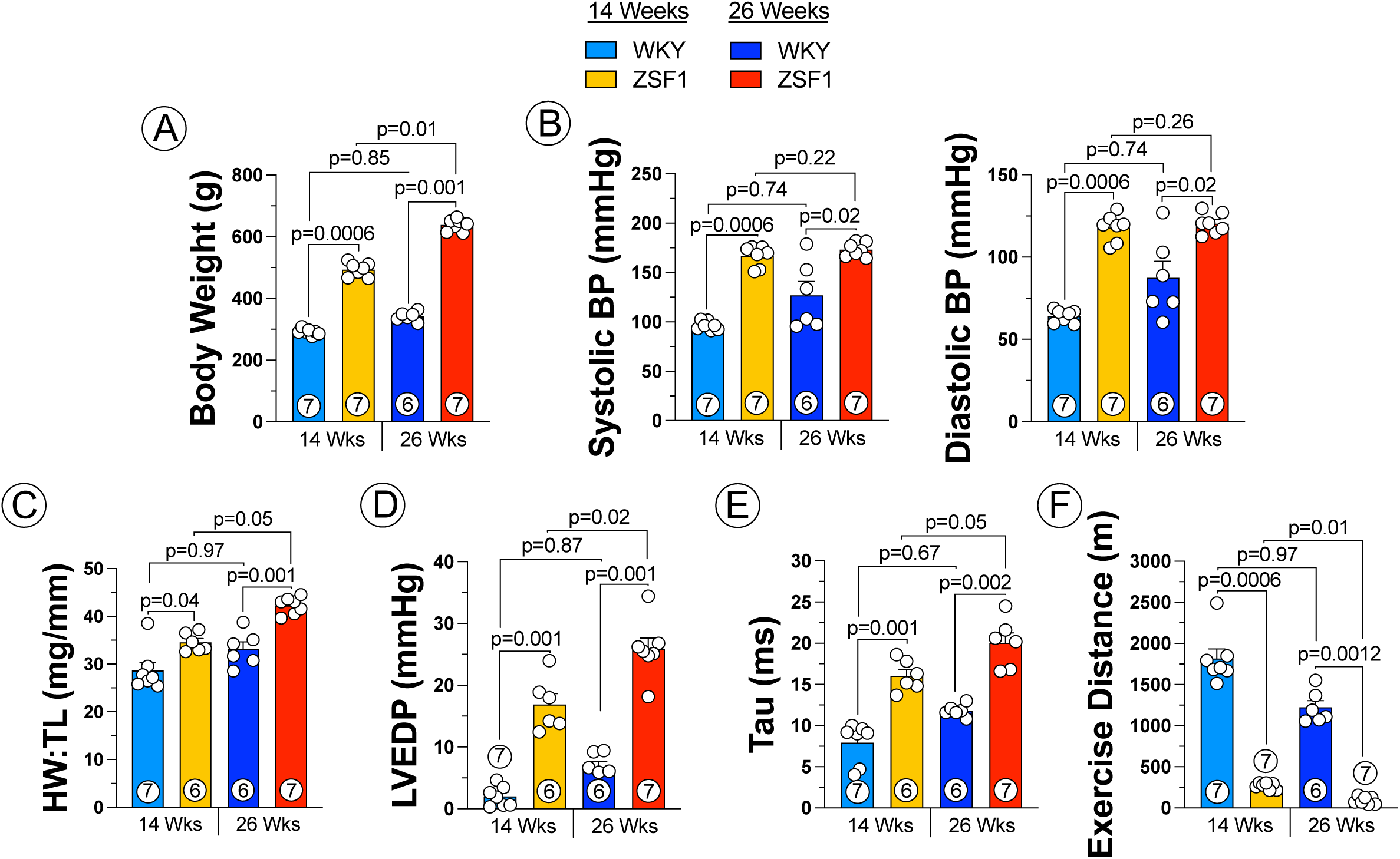
Severe Cardiometabolic HFpEF in ZSF1 Ob Rat Model of HFpEF. Body weight *(**A**)*, systolic and diastolic blood pressure *(**B**)*, heart weight over tibia length ratio (***C***), left ventricular end diastolic pressure (LVEDP) *(**D**)*, relaxation constant Tau *(**E**)*, and exercise performance in treadmill running distance *(**F**)* in WKY control vs. ZSF1 obese rats. Data were analyzed with 2-way ANOVA and presented as mean ± SEM. Circled number inside bars indicate sample size.

**Supplemental Figure 2.**
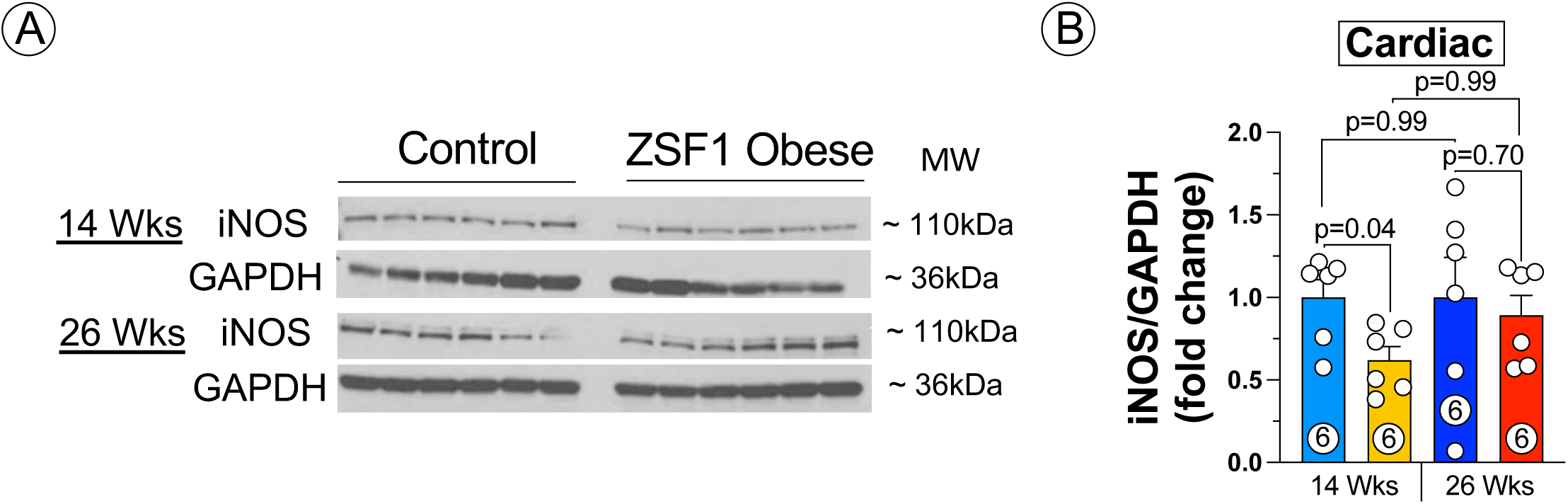
Unaltered cardiac iNOS protein expression in ZSF1 Ob Rat Model of HFpEF. Representative Western blot *(**A**)* and protein expression quantification *(**B**)* of iNOS in cardiac tissue from WKY controls vs. ZSF1 obese rats. Data were analyzed with 2-way ANOVA and presented as mean ± SEM. Circled number inside bars indicate sample size.

**Supplemental Figure 3.**
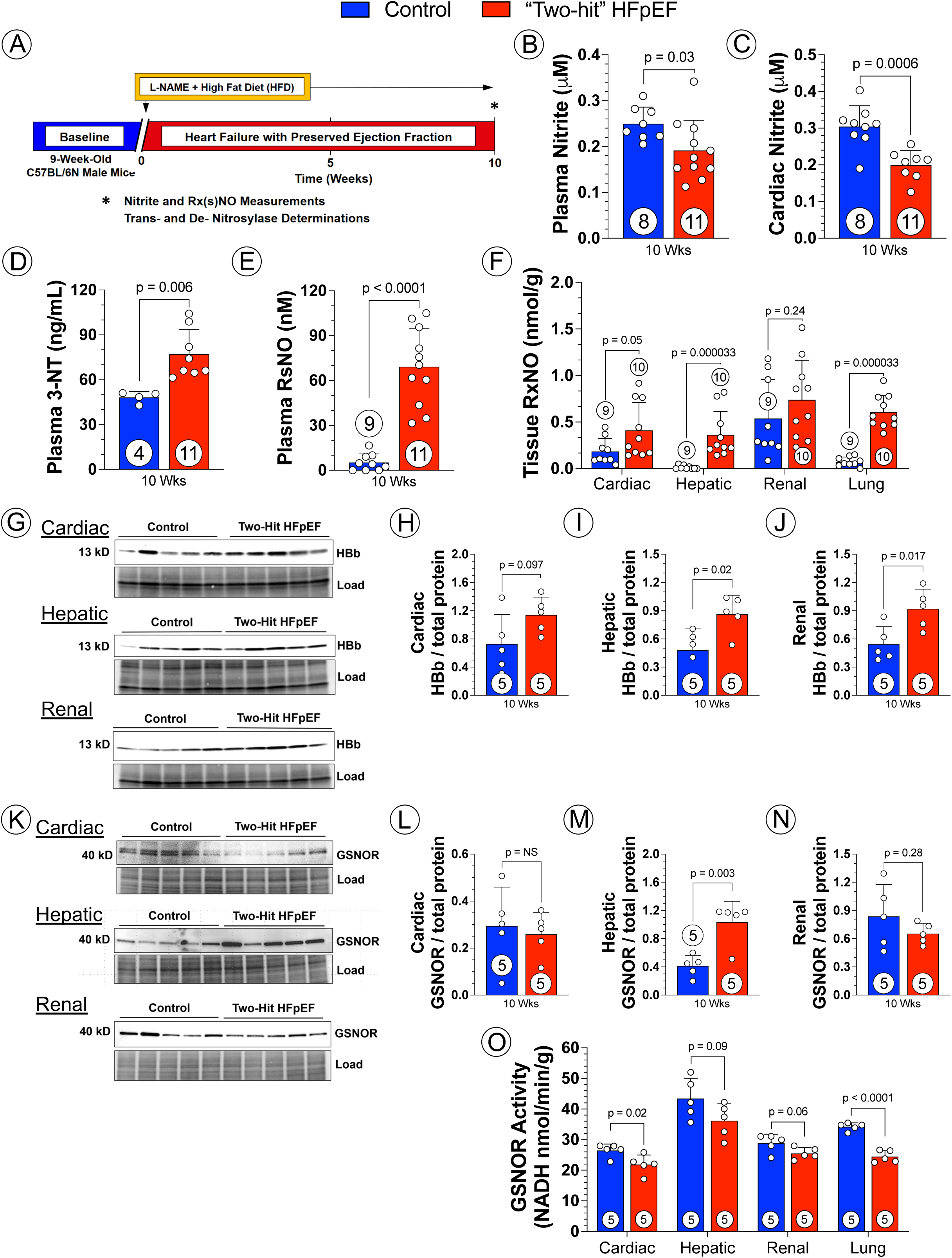
NO/RxNO Imbalance and Dysregulated Trans- and De-nitrosylases in the “two-hit” Murine Model of HFpEF. Study timeline of L-NAME + HFD “two-hit” murine model of HFpEF *(**A**)*. Circulating *(**B**)*, cardiac *(**C**)* nitrite levels from control vs. “two-hit” HFpEF mice following 10 weeks of two-hit treatment. Circulating 3-NT (***D***) and RsNO (***E***) levels control vs. “two-hit” HFpEF mice. RxNO levels in a variety of tissue including the heart, liver, kidney and lung *(**F**)* in control vs. “two-hit” HFpEF mice following 10 weeks of two-hit treatment. Representative western blot images of HBb in the heart, liver, and kidney (***G***) as well as the quantification (***H - J***) in control vs. “two-hit” HFpEF mice. Representative western blot images of GSNOR in the heart, liver, and kidney (***K***) as well as the quantification (***L - N***) in control vs. “two-hit” HFpEF mice. GSNOR enzymatic activity (***O***) in heart, liver, kidney, and lung tissue from control vs. “two-hit” HFpEF mice. Data from *(**H** - **J**)* and *(**L - O**)* were analyzed with Mann Whitney test while the other data were analyzed with student unpaired 2-tailed t-test. Data were presented as mean ± SEM. Circled number inside bars indicate sample size.

